# No support for intra-nor inter-locus sexual conflict over mating latency and copulation duration in a polyandrous fruit fly

**DOI:** 10.1101/2020.10.18.344390

**Authors:** Julie M. Collet, Jacqueline L Sztepanacz

## Abstract

The total strength of sexual selection on males depends on the relationship between various components of pre- and post-copulatory fitness. Misalignment between male and female interests creates inter-locus sexual conflict, where the fitness of one sex is increased at the expense of the other. Although rarely considered, mating behaviours can also be genetically correlated between males and females, creating intra-locus sexual conflict, where beneficial alleles in one sex are costly when expressed in the other sex. How inter- and intra-locus sexual conflicts operate on the expression of mating behaviours remains little understood. Here, we study male attractiveness, mating latency and copulation duration in two populations of the polyandrous *Drosophila serrata*. Univariate analyses show little genetic variance in mating latency, and that males, but not females, contribute to copulation duration genetic variance. Further, multivariate analyses revealed little covariance between the studied traits. However, analyses considering male and female contribution in a single framework supported genetic contributions from both sexes for mating behaviours and complex patterns of between sexes correlations. Finally, our study did not find any association between those mating behaviours and fitness component, specifically (i) no phenotypic covariance between male attractiveness and mating latency and, (ii) longer copulations did not result in the production of more offspring. With no detectable fitness benefits in any sexes for shorter mating latency or longer copulation duration, our results do not support the presence of inter-nor intra-locus sexual conflict for these mating traits.

## Introduction

The total strength of sexual selection on males depends on the relationship between various components of pre- and post-copulatory fitness (Collet et al., 2014). Theoretical expectations of how pre- and post-copulatory fitness covary, however, are unsettled. On the one hand, the phenotype-linked fertility hypothesis predicts that male ornaments reflect their fertility, and thus male pre- and post-copulatory success are predicted to be positively correlated (Sheldon, 1994). On the other, trade-offs may exist between pre- and post-copulatory traits, for example, mating success and sperm competitiveness, resulting in a negative correlation between pre- and postcopulatory fitness (Parker & Pizzari, 2010). The empirical data are also mixed: substantial experimental and observational studies support both negative (e.g. Danielsson, 2001) and positive (e.g. Collet et al., 2012) phenotypic correlations (See metanalysis Mautz et al., 2013). Despite the interest in, and the large number of phenotypic studies, there have been fewer to focus on the genetic covariances between pre- and post-copulatory traits (but see Taylor et al., 2013, Hall et al., 2013). Genetic covariances, however, are the currency for evolution, and in order to understand the evolution of copulatory traits in males, we must understand how they genetically covary.

In addition to within-sex genetic covariances, correlations of homologous traits between the sexes have an important role in determining the evolutionary trajectories of fitness-related and other traits (e.g. Gosden et al., 2012, McGlothlin et al., 2019, Sztepanacz & Houle, 2019, Holman & Jacomb, 2017). When the largely shared genome of males and females is subject to divergent selection to sex-specific phenotypes, intralocus sexual conflict can arise (Bonduriansky & Chenoweth, 2009). For example, when the alleles that underlie traits which increase male reproductive fitness, such as attractiveness, come at a cost when they are expressed in females, and vice versa. Sexual antagonism can also arise when selection is sexually concordant, suggesting that it is an inevitable by-product of species with separate sexes (Connallon & Clark, 2014). Intralocus sexual conflict results in a reduction of population mean fitness because neither sex can reach their fitness optima. Although sexual conflict may be resolved by the evolution of sexual dimorphism (Collet et al., 2016, Bonduriansky & Chenoweth, 2009), the pervasiveness of negative intersexual genetic correlations for fitness indicate that these conflicts are often unresolved (Chippindale et al., 2001, Brommer et al., 2007, Foerster et al., 2007, Kohorn, 1994, Cox & Calsbeek, 2009).

Another form of sexual conflict, interlocus sexual conflict, can arise from the interactions between the sexes, which increase the fitness of one sex at the expense of the other (Arnqvist & Rowe, 2005). These interactions result in a coevolutionary arms race where adaptation of a trait in one sex results in counteradaptation of the interacting trait in the other sex (Parker, 1979, Moore & Pizzari, 2005, Rowe et al., 2005, Arnqvist & Rowe, 2005, Chapman et al., 2003, Clutton-Brock & Parker, 1995). Such evolutionary arms races can lead to the rapid evolution of interacting traits. Historically, intra- and inter-locus sexual conflict have been treated as separate evolutionary processes. More recently, however, the potential for an interaction between intra- and inter-locus sexual conflict has been highlighted (Pennell & Morrow, 2013). In particular, theory has shown that when reproductive traits are involved in both intra- and inter-locus sexual conflict their interactions can lead males and females into a repeating cycle of conflict escalation followed by resolution, preventing them from reaching a stable equilibrium (Pennell et al., 2016).

The extent of between-sex pleiotropy (intra-locus) for traits that are also involved in inter-locus sexual conflict is scarce, and in general, we lack a comprehensive study of traits that may experience both intra- and inter-locus sexual conflict in a single framework. Copulatory traits are one class of traits that may be genetically correlated between the sexes and also interact during mating, leading to the potential for both intra- and interlocus sexual conflict. Here, we used a quantitative genetic breeding design to investigate the within and between sex heritabilities and correlations of copulatory traits in *Drosophila serrata*. Specifically, we estimated the heritability of male attractiveness, male and female mating latency and copulation duration, and the within and across-sex genetic correlations of these traits. Furthermore, we investigated whether any of these traits predicted male and female fitness, indicating the potential for sexual conflict.

The relationship between attractiveness and mating latency has been particularly well described in *Drosophila melanogaster*, where males that carry the cuticular hydrocarbon (CHC) pheromone, 7-tricosene, are more attractive (as measured by their mating rate), and mate faster, than males that do not (Grillet et al., 2006). In one study, variation in mating latency was explained to a similar extent by the genetic identity of both males and females, suggesting a role of both sexes in determining this phenotype (Tennant et al., 2014). The relationship between male attractiveness and mating latency is not universal, however. In *D. bunnanda*, males that were artificially selected to carry a more attractive combination of cuticular hydorcarbons (as measured by their mating rate) did not mate faster than males selected for a less attractive combination (McGuigan et al., 2008), and in *D. melanogaster*, MacKay *et al* (2005) found that the genetic background of females was the best predictor of mating latency. In *D. serrata*, male attractiveness has also been well described, where females have been shown to prefer a particular combination of CHC contact pheromones in males (Chung et al., 2014, Hine et al., 2002, Hine et al., 2011, Chenoweth & Blows, 2003). In a number of studies, these CHCs have consistently predicted male mating success, explaining up to 46% of its variance (Rundle et al., 2005). CHCs also genetically covary between the sexes, and may experience intralocus sexual conflict (Gosden et al., 2012). The relationship between male CHCs and mating latency, however, has not been established in this species.

In many insects, once mating starts, a cocktail of proteins (Seminal fluid proteins, Sfps) are transferred along with sperm during copulation (Poiani, 2006). These proteins trigger an array of behavioural and physiological responses in females which increase male fitness, for example, decreasing female receptivity to subsequent mating, stimulating egg laying, facilitating sperm storage, forming a mating plug, and displacing sperm from other males (Chen et al., 1988, Gioti et al., 2012, Chapman & Davies, 2004). These responses come at a cost to females, as Sfps reduce female lifespan and their lifelong reproductive success (Wigby & Chapman, 2005, Fowler & Partridge, 1989). Mating in *D. melanogaster* takes ~20 minutes, twice the time needed to transfer sperm alone (Gilchrist & Partridge, 2000), and mating duration has a genetic basis (Moehring & Mackay, 2004), and is phenotypically plastic depending on the social environment (Rouse et al., 2018). In a rare study investigating sex-specific genetic contributions to copulation duration, Edward at al. (2014) found significant genetic variance for copulation duration in both males and females *D. melanogaster*, although male’s contribution was higher than female’s.

Copulations in *D. serrata* are markedly different. In a modest study of 31 individuals, median mating duration was 4 minutes, with 157s recorded as the shortest mating duration that could lead to progeny (Hoikkala et al., 2000). Males in this species appear to transfer sperm in a single clump (J Sztepanacz pers observation), suggesting that longer copulation durations would not equate to more sperm transferred. Whether males also transfer seminal fluid proteins that affect female physiology or behaviours is unknown. Hoikkala et al (2000) observed a weak and non-significant negative relationship between copulation duration and the number of progeny produced by females, suggesting that Sfps may not have a large role in this species.

In this study we estimated the phenotypic and genetic correlations between male attractiveness, mating latency, duration, and productivity, in two independent data sets from the samespecies. Each data set was a large quantitative genetic breeding design with 80 sires and phenotypes on over 3,000 flies measured in each experiment. These independently replicated data provide unprecedented power to identify the genetic basis of copulatory behaviours and their effects on fitness.

## Materials and methods

### Experimental and Breeding Design

The data that we analyse here come from two independent populations and experiments in *D. serrata* performed under similar conditions. The data from Population 1 allows us to estimate the heritabilities for and genetic correlations between male attractiveness, mating latency, and mating duration. The data from Population 2 allows us to estimate the phenotypic correlations between male attractiveness and mating latency, and heritabilities for and genetic correlations between male and female mating latency and duration, and their cross-sex genetic correlations. Further, this experiment measured productivity of the resulting matings, allowing us to directly estimate the relationship between mating behaviour and fitness. Therefore, the combination of these two experiments enable us to track the outcomes of mating interactions from pre- to post-copulatory fitness.

#### Population 1

Experiments in population 1 were conducted on an outbred laboratory population of *D. serrata* (Rundle et al., 2006) maintained at a large population size (N>2000) under standard laboratory conditions. A half sibling breeding design was carried out to estimate additive genetic variances and covariances in traits of interest. The details of the breeding design are described in detail in Sztepanacz and Rundle (2012). Briefly, eighty sires were each mated to three virgin dams, which were allowed to oviposit for 72h after mating. Upon emergence of their offspring, male offspring were collected from these families using CO2 anaesthesia and were held as virgins at a density of 6 flies per vial for 5-7 days prior to their use in experimental assays. The breeding design was conducted in two blocks of 40 sires, spanning two generations of the laboratory population, and resulted in 2941 individuals from 240 full- and half-sibling families from 80 sires. Brothers from each family were used in either a (1) a competitive mating trial where their cuticular hydrocarbons (CHCs) were subsequently extracted or (2) in a behavioural assay which measured mating latency and duration.

The details of the binomial mate choice and CHC assay are described in detail in Sztepanacz and Rundle (2012), where these data were first published. They found that male CHCs were under significant directional sexual selection which explained 9.1% of the variance in male mating success. In order to obtain an ‘attractiveness’ score for these males, here we applied the standardized sexual selection gradient *β*, presented in their paper, to the individual pheromone profiles of each male (n=1979 males).

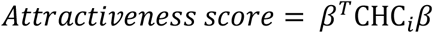

Where *β* is the standardized sexual selection gradient, ^⊤^ denotes the transpose of the vector, and CHC_i_ indicates the vector of 8 CHC traits for an individual.

To measure mating latency and duration a single male fly was introduced in a vial with a single female which had the same genetic background as the male, but was fixed for a recessive orange-eye mutation. The time it took before mating began was recorded as mating latency. Once mating started, copulation duration was recorded as the time during which genital contact could be observed (n=962 males). If the pair did not mate during the first two hours, the pair was discarded from the measure of mating latency and copulation duration.

#### Population 2

Experiments in population 2 were conducted on an outbred laboratory population of *D. serrata* described elsewhere (Hine et al., 2014), maintained with a large population (N>2000) under standard laboratory conditions. First, we carried out a paternal half-sibling breeding design to estimate the genetic variances and covariances between mating latency, duration, and productivity in males and females. Eighty sires were each mated with three females and up to 10 female and 10 male virgin offspring of each pair were collected using ice anaesthesia. Males and females were kept separately (up to four males per vial, up to six females per vial) in vials containing 7mL of standard food, for four to six days before mating trials. The day before the experiment, males and females from the breeding design were transferred in an individual vial with fresh food. In total 1558 males and 1726 females, were produced across three blocks from 80 sires and 220 dams. Males and females from each family were used in (1) behavioural assays to measure mating latency and duration, and (2) had their productivity subsequent measured.

On the day of the experiment, each focal female was put together with a virgin male from the same population and time until mating started was recorded as mating latency. If the pair did not mate during the first two hours, the pair was discarded from the measure of mating latency and copulation duration.

Once mating started, copulation duration was recorded as the time during which genitalia contact could be observed. At the end of the mating, males were discarded, and females placed in individual 10mL vials with standard food and without live yeast. We left females lay eggs for 24 hours, after which females were discarded and offspring were left to develop. We chose 24h to capture the potential male effect of mating on the number of eggs laid, that was shown to occur when sex peptide was injected into female *D. melanogaster* (Aigaki et al., 1991). After offspring emergence, vials with adult offspring were frozen at −20C and the number of adults that had emerged was counted as the measure of productivity.

Males from the paternal half-sibs were tested in similar conditions. The day before mating trial, males from the breeding design were isolated in a fresh vial with 10mL standard food. On the mating trial day, each male was put together with a virgin female that was fed with *ad libitum* yeast. Mating latency and copulation duration were recorded the same way as for females. Male productivity was also recorded as the number of offspring that emerged from the vial of their female mate.

When recording productivity from the frozen vials, there was a few instances where the number of offspring was impossible to individually count, as they emerged in the stopper. In a few vials, we proceeded to careful visual inspection to estimate that around five flies must have been caught in the stopper. Thus, five additional offspring were systematically added to the productivity obtained in subsequent vials in which those emergence were encountered.

Finally, we measured whether attractive males mated faster. We randomly collected virgin males and females at emergence from the population. Four to six days after collection, to guarantee sexual maturity, two virgin males were put together with a virgin female. The time between the introduction of the female in the vial with both males and the beginning of the mating was recorded as mating latency. After copulation started, the mating pair was separated and the male who mated (chosen) and the other male (rejected) had their CHCs extracted and assayed using gas chromatography following standard procedure as in population 1 (Blows et al., 2004, Sztepanacz & Rundle, 2012). We then followed the same method as in population 1 to obtain attractiveness scores; we determined the selection gradients for male CHC profile by using the partial regression coefficients of the linear regression of standardized mating success on the standardized log contrasts of CHCs peaks, and applied it to the individual pheromone profile of each male.

### Statistical analyses

#### Quantitative genetic analyses in Populations 1 and 2: heritabilities and genetic covariances

All models estimating quantitative genetic parameters in both populations were performed on standardized (z-score) data using animal models (Lynch & Walsh, 1998, De Villemereuil, 2012) with the R package MCMCglmm (Hadfield, 2010). In all models, blocks were entered as fixed effects and additive genetic effects (using the pedigree information) were entered as random effects. In models that included attractiveness in population 1, a supplementary *column* effect was added to account for the use of two different gas chromatography columns for this experiment. Error distributions were Gaussian. Number of iterations, burn in, and thinning intervals varied for each model, and they were set to achieve convergence by using the Heidelberger and Welch test in the coda R package (Heidelberger & Welch, 1981, Plummer et al., 2006) and a minimum effective sample of 1000 for all studied parameters. All credible intervals are 89% (Makowski et al., 2019). Analyses of Population 1 and Population 2 were performed independently.

First, univariate animal mixed models were performed on mating latency, copulation duration, and attractiveness, in females and in males. Prior distributions were sets to inverse-Gamma, with parameters V=1 and nu=0.002. Heritabilities were calculated from the posterior distribution as the median proportion of phenotypic variance explained by the animal factor. To confirm whether there was additive genetic variance, we compared the Deviance Information Criterion (DIC) of these univariate models to the DIC of models that did not include the ‘animal’ random effect. DIC are useful to select the model which best describes the data (as for example the AIC), when the posterior distribution is well summarized by its mean (Spiegelhalter et al., 2002, Gelman et al., 2014).

To investigate covariance between pre- and post-copulatory traits, and between the sexes, we ran multivariate animal mixed models including the traits for which we tested genetic covariance with MCMCglmm. Additive genetic variance-covariance G matrices were inferred from the median posterior distribution of genetic variances and covariances of the animal effect. To test the extent of genetic covariance, we compared the DIC of those models to the DIC of models in which the genetic covariance was set to 0 (*idh*, off-diagonal of the G matrix =0).

#### Phenotypic analyses in population 2: relationships between copulatory and fitness traits

We used population 2 to investigate whether mating latency or copulation duration could predict the measured components of pre- and post-copulatory fitness. First, we used the mate choice experiment to see whether more attractive males mated faster. We tested whether the log transformed mating latency could predict the attractiveness score with a linear model (lm in R-Core-Team, 2014) in males that were successful at mating.

Further, we tested whether mating latency or copulation duration predicted the pair’s productivity. Because the dataset was obtained in the paternal half-sibs breeding design, we accounted for relatedness between tested individuals with the following mixed model (Pinheiro et al., 2020):

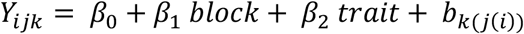

where *Y*_*ijk*_ is the standardized productivity of the *k*^th^ male, son of the *j*^th^ dam and *i*^th^ sire, *block* is to account for the three blocks performed, *trait* either mating latency or copulation duration.*b*_*ijk*_ is an observation (*k*)-level random effects nested in dams (*j*), themselves nested in sires (*i*). Statistical significance for the *trait* effect was tested with Log-Likelihood ratio tests comparing two models where the *trait* fixed effect was included or not.

## Results

### Heritability of attractiveness, mating latency, and mating duration

There was substantial phenotypic variation in male attractiveness scores in both populations, which had a heritability of 0.31 in population 1 (estimated using REML, genetic effect vs. no genetic effect ΔAIC=75.2, Suppl Materials, MCMCglmm ΔDIC=1018, Table 1). Mating occurred shortly after males and females were put together in the vial. Over a quarter of the pairs mated in the first 5 minutes in Population 1 and in the first 7 minutes in Population 2, however the distribution was skewed with a long tail extending to 120min, which is when the experiment was stopped (Fig. 1). The median mating latency in Population 1 was 10.2 minutes, and in Population 2 was 22.4 minutes (Fig. 1). We estimated the heritability of mating latency using the observed variance standardised data, and using log-transformed values. Overall, the estimates of heritability were similar, so we report the non-transformed values here and heritability of log-latency in the supplementary material (Supplementary Table 2). Univariate analyses in population 2 found no evidence for heritable variation in mating latency in males (*h^2^*=0.007; CI [0.000; 0.046]) nor in females (*h^2^*= 0.009; CI [0.001; 0.050]). In population 1, however, there was some evidence that latency was heritable in males (*h^2^*= 0.068; CI [0.003; 0.175]) and evolvable (*e* = 1.3%). In *D. melanogaster*, single phenotypic observations of latency have been shown to be a highly noisy measure (Hoffmann, 1999), which may explain why we were able to detect heritable variation in one population but not the other.

**TABLE 1:** Summaries of univariate models. h^2^ is the median narrow sense heritability, Eff. Samp. is the Effective Sample Size, DIC dif is thedifference in the deviance Information Criterion between the null model and the tested models. Credible intervals are 89 %. *models for those estimates did not converged according to the Heidelberger and Welch test.

| Population | Parameter | $h^2$ | Credible interval | Eff. Samp. | DIC dif | evolvability (%) | Credible interval |
| --- | --- | --- | --- | --- | --- | --- | --- |
| 1 | Male Latency | 0.068 | [0.003 ; 0.175] | 1871 | 10 | 1.3 | [0.0 ; 26.0] |
|  | Male Duration | 0.153 | [0.064 ; 0.257] | 2737 | 35 | 1.5 | [0.7 ; 2.6] |
|  | Male attractiveness | 0.678 | [0.576 ; 0.787] | 26397 | 1018 |  |  |
| 2 | Male Latency | 0.007 | [0.000 ; 0.046] | 1138 | -1 | 0.0* | [0.0 ; 2.1]* |
|  | Male Duration | 0.043 | [0.002 ; 0.043] | 1135 | 6 | 0.2 | [0.0 ; 0.6] |
|  | Female Latency | 0.009 | [0.001 ; 0.050] | 1724 | 0 | 0.0* | [0.0 ; 2.5]* |
|  | Female Duration | 0.005 | [0.000 ; 0.031] | 1084 | -1 | 0 | [0.0 ; 0.2] |

**Figure 1:**
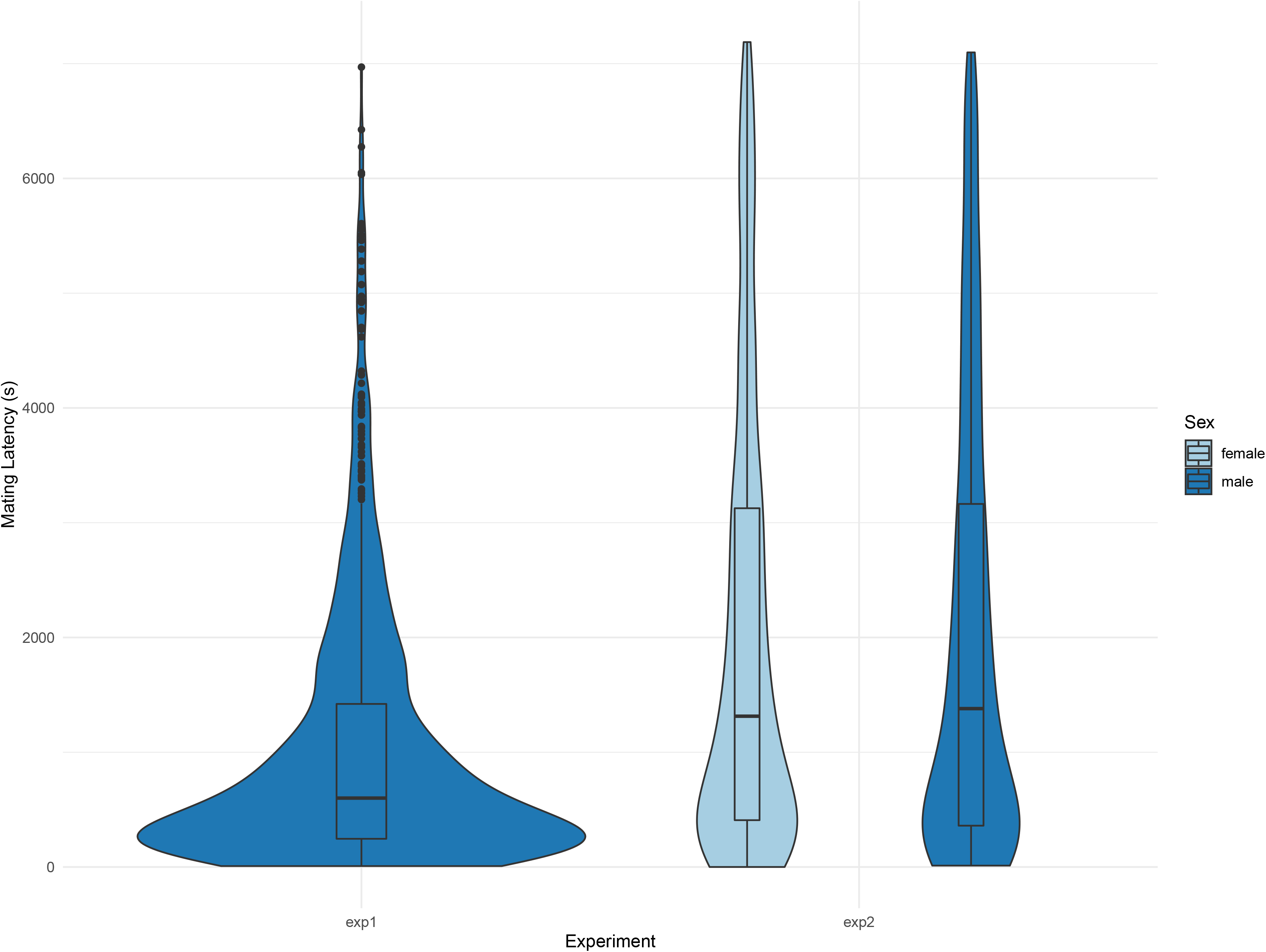
Distribution of mating latencies (in seconds) in population 1 and population 2. In population 2, results are shown in two different colours according to whether they were measured in females or males of the breeding design.

Once mating started, the median copulation duration in Population 1 was 5.8 minutes and 4.6 minutes in Population 2 (Fig. 2). This is consistent with the durations observed in *D. serrata* by Hoikkala et al (2000), which ranged from 2.62 to 7.87 minutes with a median of 4 minutes. Copulation duration was heritable in males from Population 1 with an estimated heritability of 0.15 and an evolvability of 1.5%, suggesting that this trait has a genetic basis and can respond to selection (Table 1). In Population 2, however, heritabilities were low in both sexes, although higher in males (male *h^2^* = 0.043 [0.002; 0.043]; female *h^2^* = 0.005 [0.000; 0.031]), with little statistical support (male ΔDIC = 6; female ΔDIC = −1, Table 1).

**Figure 2:**
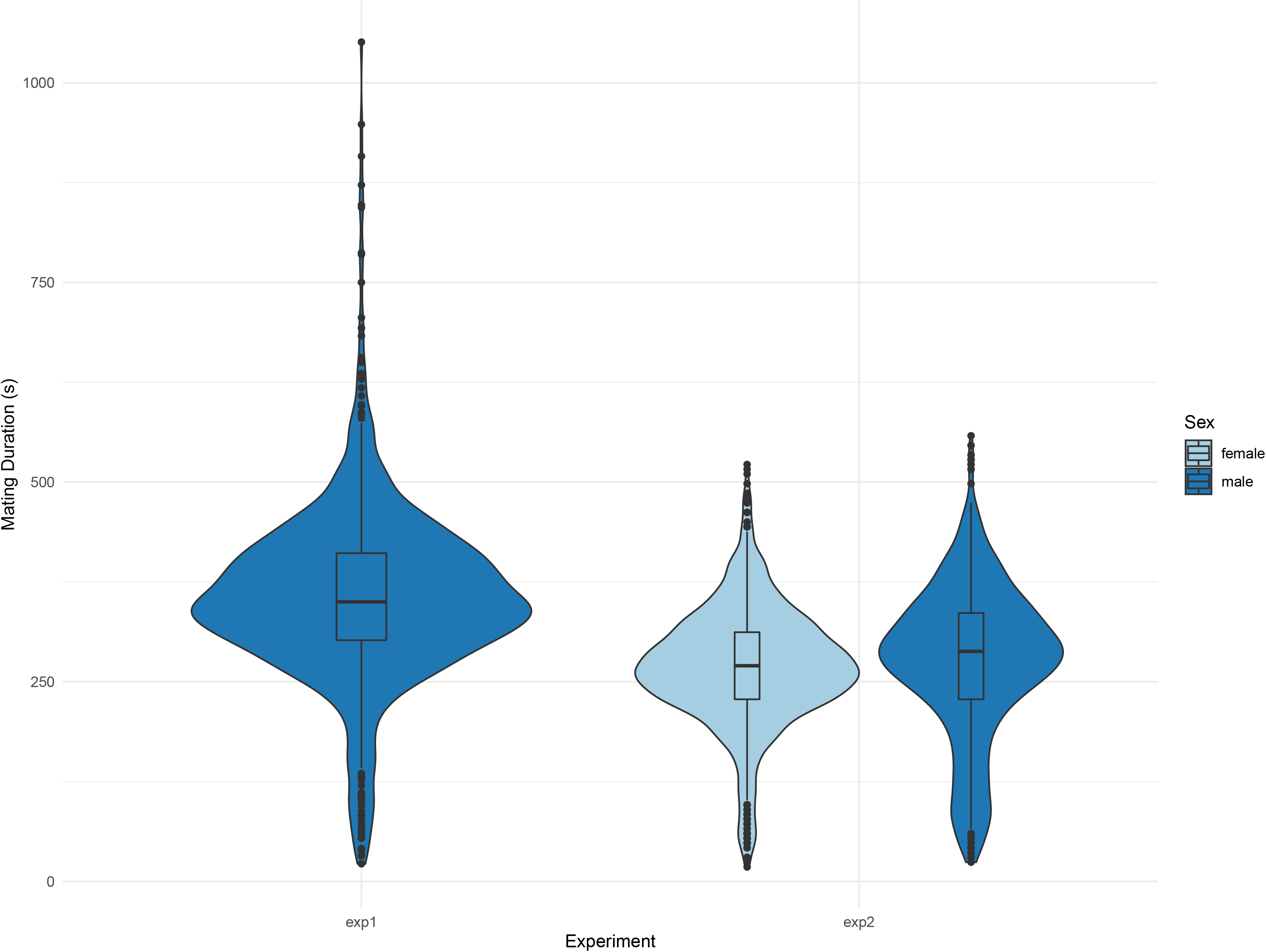
Productivity of mating pairs according to the duration of their copulation. Darker colours appear when several datapoints overlap. The line corresponds to a regression line obtained with the mixed models accounting for the individuals relatedness.

### Relationships between copulatory traits

There was no genetic correlation between male attractiveness score and mating latency, with a credible interval that overlapped zero (Bivariate model in Population 1: *r_G:_* 0.11; [−0.17; 0.37]; ΔDIC = 21, Table 2). The genetic covariance matrix for male attractiveness, mating latency, and mating duration in Population 1 is shown in Table 2. Overall, the genetic correlations were low between all traits with confidence intervals of the estimates overlapping 0 and ΔDIC between models with and without genetic covariance equal to −2.

**TABLE 2:** The genetic variance-covariance matrix, G for male attractiveness, mating latency, and copulation duration in population 1. Heritabilities are in bold along the diagonal, covariances are below the diagonal, and correlations are in italic above the diagonal. 89% confidence interval for each value are in brackets.

|  | Attractiveness | Latency | Duration |
| --- | --- | --- | --- |
| Attractiveness | **0.636 [0.553 ; 0.716]** | *0.102 [-0.152 ; 0.347]* | *-0.059 [-0.281 ; 0.176]* |
| Latency | 0.009 [-0.015 ; 0.035] | **0.133 [0.076 ; 0.217]** | *-0.062 [-0.421 ; 0.316]* |
| Duration | -0.006 [-0.032 ; 0.020] | -0.009 [-0.071 ; 0.050] | **0.179 [0.104 ; 0.276]** |

### Cross-sex relationships for interactive traits

In population 2 we were able to estimate both within-sex and across-sex genetic covariances for mating latency and duration. The full genetic correlation matrix is shown in Table 3. Similar to population 1, genetic correlations between latency and duration within each sex were modest with credible intervals of the estimates overlapping 0. However, the overall G matrix contained some level of genetic covariance, as ΔDIC between models with and without genetic covariance equal to 95.

**TABLE 3:** The genetic variance-covariance matrix, G_*f*_*m* for latency and mating duration in population 2. Heritabilities are in bold along the diagonal, covariances are below the diagonal, and correlations are in italic above the diagonal. 89% confidence interval for each value are in brackets.

|  |  | Female |  | Male |  |
| --- | --- | --- | --- | --- | --- |
|  |  | Latency | Duration | Latency | Duration |
| Female | Latency | **0.084 [0.051 ; 0.133]** | *-0.081 [-0.392 ; 0.251]* | *0.111 [-0.239 ; 0.431]* | *0.094 [-0.260 ; 0.416]* |
|  | Duration | -0.006 [-0.033 ; 0.020] | **0.070 [0.044 ; 0.108]** | *-0.056 [-0.379 ; 0.283]* | *-0.170 [-0.473 ; 0.171]* |
| Male | Latency | 0.010 [-0.021 ; 0.043] | -0.004 [-0.033 ; 0.023] | **0.087 [0.052 ; 0.142]** | *-0.126 [-0.456 ; 0.239]* |
|  | Duration | 0.009 [-0.026 ; 0.045] | -0.014 [-0.049 ; 0.015] | -0.012 [-0.053 ; 0.024] | **0.116 [0.069 ; 0.183]** |

Consistent with other studies that estimate cross-sex genetic covariance matrices (Gosden & Chenoweth, 2014, Sztepanacz & Houle, 2019), we found that estimates of cross-sex cross-trait correlations were asymmetric. The point estimate of the correlation between latency in males and duration in females was negative (*r*_*G*_ = −0.056; [−0.379; 0.283]), while the correlation between latency in females and duration in males was positive (*r*_*G*_ = 0.094; [−0.260; 0.416]). The confidence intervals of these estimates were large and overlapping however, so we cannot say with confidence that they differ from each other

### Relationships between copulatory traits and fitness components

In population 2 we were able to determine the relationship between mating latency, copulation duration, and two fitness components: male attractiveness (determined from mating success assays) and productivity. There was no phenotypic relationship between male attractiveness and mating latency (Population 2: F_1,199_ = 0.83, p=0.36, Fig. 3A) nor with the pair productivity (F_1,1781_ = 0.14, p=0.71, Fig. 3B). Finally, copulation duration showed a tendency to be negatively associated with the pair productivity (F_1,1754_=3.6, p=0.06, Fig. 4).

**Figure 3:**
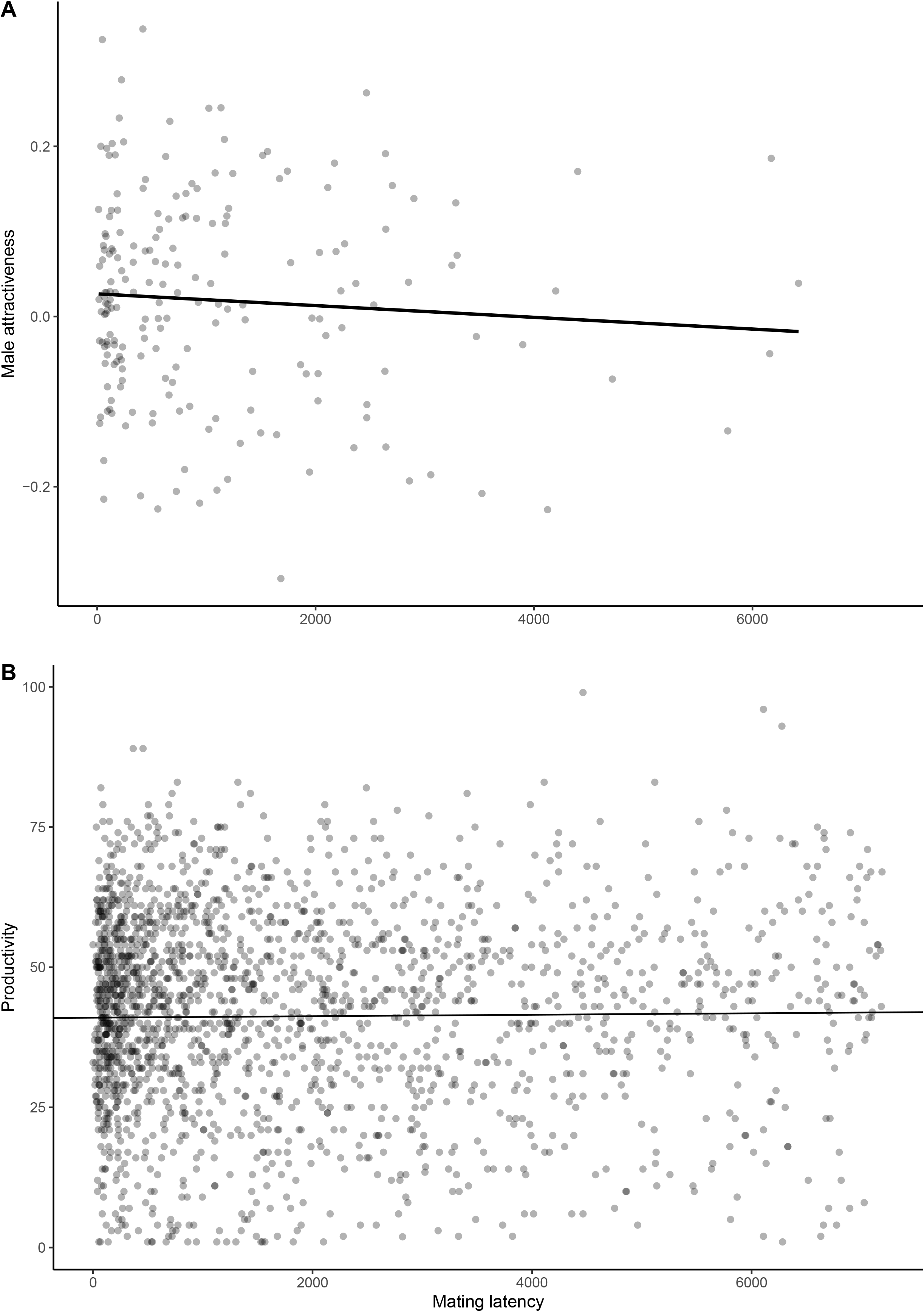
Mating latency (in seconds) as a predictor of male fitness: either **A:** male attractiveness or **B:** productivity of a pair. **A**: Each datapoint represents a successful male’s latency from introduction into a vial with a virgin female to copulation, and their attractiveness index based on their CHCs profiles. The line corresponds to a linear model (“lm” in R). **B**: Each datapoint represents a mating pair. Darker colours appear when several datapoints overlap. The line corresponds to the regression line obtained using a mixed model accounting for the replication due to relatedness between individuals.

**Figure 4:**
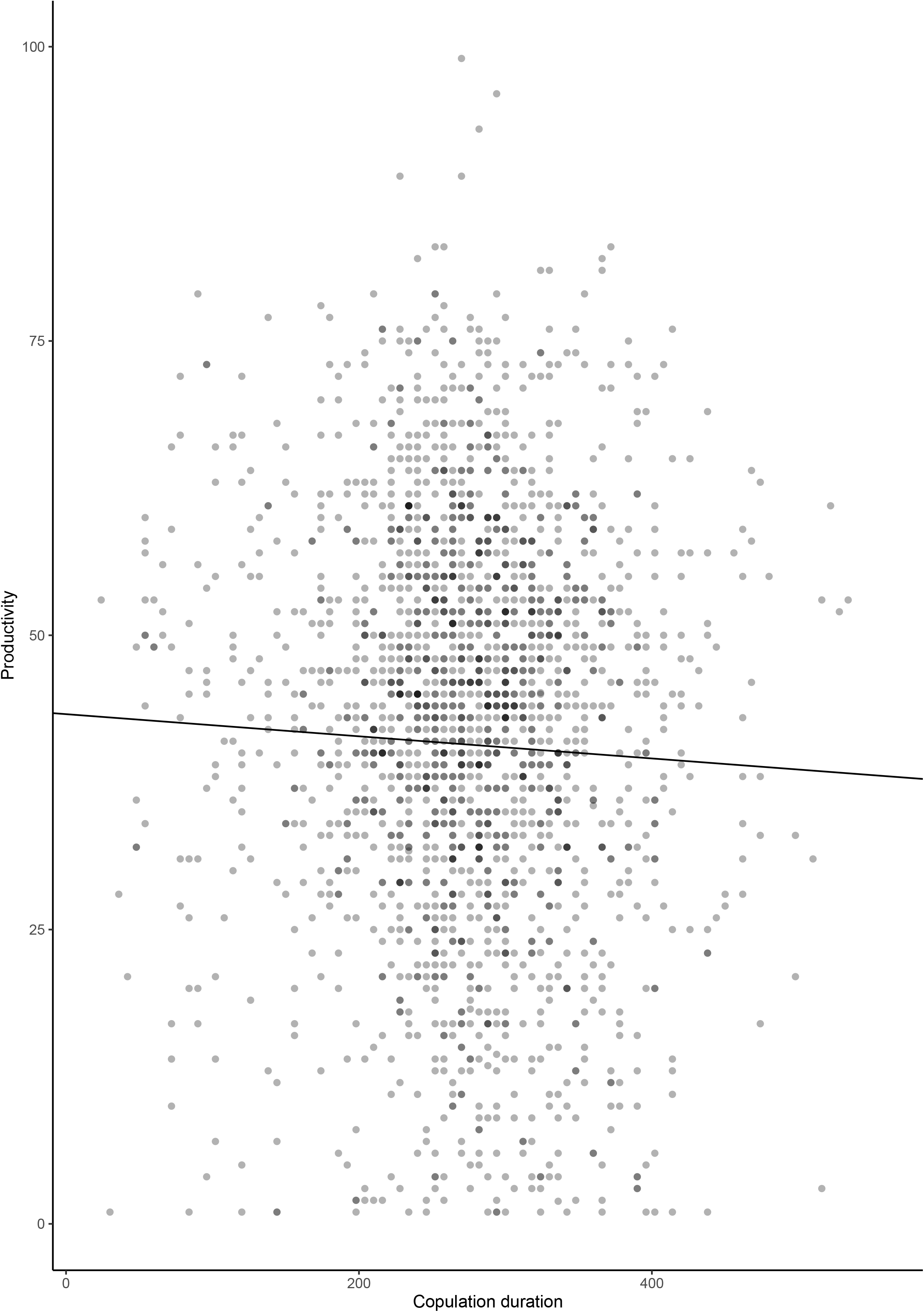
Distribution of copulation durations (in units of seconds) in population 1 and population 2. In population 2, results are shown in two different colours according to whether they were measured in females or males of the breeding design.

## Discussion

This study provides a unique overview of the fitness effects and genetic contributions of two widely studied interactive traits, mating latency and copulation duration, in a species with higher polyandry and shorter copulations than traditional *Drosophila* model systems. We found that, in *Drosophila serrata*, (i) copulation duration was heritable in males in both population, (ii) multivariate analyses provided more power to detect male and female contributions to interactive traits, and (iii) mating latency and copulation duration did not predict the two measured fitness components, namely male attractiveness and short-term productivity.

Quantitative genetic analyses showed that variation in copulation duration was heritable in males from both populations. When only one of both sexes genetically contribute to an interactive trait, it limits the opportunity for arm race, as adaptation would only respond to selection on one of the sexes (although plastic behaviours may still enable conflictual interactions, Moore & Pizzari, 2005). However, female genetic contribution to copulation duration is significantly higher than 0, as revealed by multivariate analyses (Table 3), so the opportunity for conflict is not absent.

Male and female genetic contributions to copulation duration were very similar in this study than in *D. melanogaster* (h^2^ _female_ = 0.08 ± 0.04 in *D. melanogaster vs* h^2^ _female_ = 0.08 ± 0.05 in *D. serrata*, and h^2^ _male_ = 0.13 ± 0.05 in *D. melanogaster vs* h^2^ _male_ = 0.15 ± 0.10 and h^2^ _male_ = 0.12 ± 0.06 in population 2 of *D. serrata*). To obtain these results in *D. melanogaster*, Edward *et al.* (2014) also used a paternal half sibs, but they mated pairs that both came from the breeding design in a full-factorial design. Thus, their multivariate model incorporated both male and female genetic contributions, as did our second, multivariate model. Sex-specific genetic contributions to copulation duration may thus be conserved between species.

The differences in heritability estimates between univariate and multivariate models that incorporated both sexes in population 2 were important, underscoring the value of incorporating both sexes’ genetic contributions in a single model when studying traits that can cause sexual conflict. However, we did not find any significant pairwise correlations between pre- and post-copulatory traits, or across sexes. Other studies of multivariate cross-sex genetic covariances between copulation behaviours are rare. Edward et al (2014), found support for within female genetic covariance for number of eggs laid before and after mating in *D. melanogaster*, whilst Han et al. (2020) found support for inter sexual genetic correlation for male mate guarding and female latency to mate in field crickets. Our comprehensive approach considering several traits in both sexes in a single framework showed that the cross-sex G matrix carried complex and statistically supported covariance between interactive traits and sexes.

Male attractiveness was not correlated with mating latency at the phenotypic or genetic level in our study. Consistent with this result, several experiments selecting for more attractive and more competitive males have failed to detect a change in their mating latency. In *D. pseudoobscura* (Bacigalupe et al., 2008), *Callosobruchus maculatus* (Maklakov et al., 2010), and even *D. melanogaster* (Nandy et al., 2013), males under male biased sex ratio, an experimental condition expected to favour attractive males, did not have a different mating latency than males evolved under equal or female biased sex ratios. In *D. bunnanda*, artificially selected attractive males also failed to mate faster (McGuigan et al., 2008), which the authors suggest was due to a lack of variation in attractiveness rather than latency being a non-heritable trait in males. Mating latency may also vary with female attractiveness, but we could not test this using our data. Mating latency could also be due to female responsiveness and choosiness, however, the heritability of mating latency in females was as low as in males. Finally, we may have failed to detect genetic contributions to our traits because of our experimental protocol. Mating traits are notoriously difficult to capture in single assays. Hoffman (1999) found that mating latency did not appear heritable after one mating trial, however when individuals were tested several times for the same mating trait it was possible to detect non-zero heritability. Although our experimental protocol created numerous repetitive measures within families (on average 28 sons tested per sire, 160 sires across populations), there was no repetition of individual measures and environmental noise of single measures may have overwhelmed our ability to detect genetic variation.

We did not find that longer matings resulted in more offspring. Indeed, our results showed a non-significant trend that longer copulations resulted in fewer offspring produced, consistent with that observed by Hoikkala et al (2000). In *D. melanogaster*, sperm transfer only takes half of the time of copulation and the second half is used to transfer seminal fluid proteins (Gilchrist & Partridge, 2000). Those proteins affect numerous female functions, in particular increase the number of eggs they lay and reduce their receptivity to subsequent mating (Chen et al., 1988, Gioti et al., 2012). In *D. serrata*, sperm are transferred in a single ball (Sztepanacz, pers. comm.), and little is known about Sfps transfer. Wing song, however, has been shown to be an important component of copulatory courtship display, with females discriminating against males that were not able to produce courtship song during copulation (Hoikkala et al 2000.). Wing shape in male *D. serrata* is genetically variable (Sztepanacz & Blows, 2015), and in other species there is weak evidence that shape variation may be associated with variation in wing-song (Menezes et al., 2013, Snook et al., 2005). Whether *D. serrata* males with different wing shapes produce different courtship songs, and whether song variation explains variation in copulation duration is unknown. *D. serrata* males have also been shown to make active courtship and mounting attempts after copulation (Hoikkala et al 2000.), which may keep the female unreceptive, preventing further matings and sperm displacement (Alcock & Buchmann, 1985). Although relatively little is known about male manipulation after mating, our results suggest that physiological effects of seminal fluid proteins may be limited in this species.

Altogether, our comprehensive analyses of mating latency and copulation duration provide little support for the hypothesis that these traits could be subject to intra- or inter-locus sexual conflict, as they were not phenotypically or genetically associated with any pre- and copulatory components of fitness studied here. We cannot rule out a lack of experimental power as a cause of low heritability estimates, however, we analysed data from over 6000 flies and 160 sires, underscoring the difficulty in estimating quantitative genetics parameters on behavioural traits. Multivariate cross-sex genetic analyses revealed that complex patterns of between sexes correlations could set the scene for complex evolution for these interactive traits.

## Acknowledgements

JMC is very grateful to Jessica McCarroll and Grace Stanton for their valuable help in the laboratory, and to Mark Blows for his excellent suggestions for the experimental design and to support this project. JLS thanks Howard Rundle for supporting this work, and D. Arbuthnott, M. Delcourt, and K. MacLellan for help in the lab. JMC’s contributions to this work was funded by the Australian Research Council, and JLS’s contributions were funded by NSERC.

