## Supplementary material for "No support for intra-nor inter-locus sexual conflict over mating latency and copulation duration in a polyandrous fruit fly"

**Supplementary Methods and Results:**

Quantitative genetic parameters were estimated using REML in the program WOMBAT (Meyer 2007), to confirm the results obtained using MCMCglmm. The equivalent models were fit with the same fixed and random effects, and confidence intervals of the point estimates were obtained using REML-MVN sampling from the inverse of the Fisher Information Matrix (Houle and Meyer 2015, Sztepanacz and Blows 2017). The confidence intervals obtained from REML-MVN implemented in WOMBAT are analogous to the credible intervals obtained from the posterior distribution of MCMCglmm samples and are subject to the same boundary constraints where variances must be $\geq 0$ (Sztepanacz and Blows 2017). For this reason, comparing the lower confidence interval of a variance estimate to 0, to ascertain whether there is non-zero genetic variation (or heritability), is inappropriate.

**Supplementary Table 1:** Variance component estimates and 90% confidence intervals of univariate models estimated using REML in WOMBAT. Confidence intervals were obtained from 1000 REML-MVN samples. REML-MVN sampling failed in two cases where there was no detectible additive genetic variation in the data.

| Population | Parameter | $h^2$ | Confidence Interval |
|---|---|---|---|
| 1 | Male Latency | 0.076 | [0.024; 0.169] |
| | Male Duration | 0.11 | [0.048; 0.208] |
| | Male Attractiveness | 0.31 | [0.237; 0.410] |
| 2 | Male Latency | 2.89E-06 | REML-MVN failed |
| | Male Duration | 0.064 | [0.022; 0.177] |
| | Female Latency | 0.011 | [0.001; 0.148] |
| | Female Duration | 1.08E-06 | REML-MVN failed |

**Supplementary Table 2:** Variance component estimates and 90% confidence intervals of univariate models of mating latency after log transformation and standardization to mean of 0 and variance of 1. Estimates were obtained using REML in WOMBAT, and confidence intervals were obtained from 1000 REML-MVN samples. REML-MVN failed in one case where there was no detectible additive genetic variation in the data.

| Population | Parameter | $h^2$ | Confidence Interval |
|---|---|---|---|
| 1 | Log Male Latency | 0.076 | [0.031; 0.193] |
| 2 | Log Male Latency | 2.89E-06 | REML-MVN failed |
| | Log Female Latency | 0.001 | [ 0.001; 0.045] |
